# TANGO2-related rhabdomyolysis symptoms are associated with abnormal autophagy functioning

**DOI:** 10.1101/2023.03.29.534583

**Authors:** Hortense de Calbiac, Sebastian Montealegre, Marjolène Straube, Solène Renault, Hugo Debruge, Loïc Chentout, Sorana Ciura, Apolline Imbard, Edouard Le Guillou, Anca Marian, Nicolas Goudin, Laure Caccavelli, Sylvie Fabrega, Arnaud Hubas, Peter van Endert, Nicolas Dupont, Julien Diana, Edor Kabashi, Pascale de Lonlay

**Affiliations:** Université Paris Cité, INSERM, CNRS, Institut Necker Enfants Malades, F-75015 Paris, France; Reference Center of Inherited Metabolic Diseases, Hôpital Universitaire Necker-Enfants Malades,, AP-HP, Institut Imagine, Filière G2M, MetabERN, F-75015, Paris, France; Translational Research for Neurological Diseases, Institut Imagine, INSERM UMR 1163, Université Paris Cité, F-75015, Paris, France; Metabolic biochemistry, Hôpital Universitaire Necker-Enfants Malades, AP-HP, Institut Imagine, Filière G2M, MetabERN, Université Paris Cité, F-75015, Paris, France; Cell Imaging & Flow Cytometry Core Facilities, Structure Fédérative de Recherche Necker, INSERM US24/CNRS UMS3633, F-75015, France; Platform, Structure Fédérative de Recherche Necker, F-75015, Paris, France; Genetics and Molecular Biology, Laboratoire de culture cellulaire, Hôpital Universitaire Cochin, AP-HP, F-75014, Paris, France; Service Immunologie Biologique, AP-HP, Hôpital Universitaire Necker-Enfants Malades, F-75015, Paris, France

**Keywords:** autophagy, calpeptin, myoblasts, rhabdomyolysis, TANGO2, zebrafish

## Abstract

Patients with pathogenic variants in the TANGO2 gene suffer from severe and recurrent rhabdomyolysis (RM) episodes precipitated by fasting. Since starvation promotes autophagy induction, we wondered whether TANGO2-related muscle symptoms result from autophagy insufficiency to meet cellular demands in stress conditions. Autophagy functioning was analyzed *in vitro*, in primary skeletal muscle cells from TANGO2 patients in basal and fasting conditions. In addition, we developed a tango2 morphant zebrafish model to assess the effect of *tango2* knockdown (KD) on locomotor function and autophagy efficiency *in vivo*. We report that TANGO2 mutations are associated with decreased LC3-II levels upon starvation in primary muscle cells, but not in fibroblasts. In zebrafish larvae, *tango2* knockdown induces locomotor defects characterized by reduced evoked movements which are exacerbated by exposure to atorvastatin, a compound known to cause RM. Importantly, RM features of *tango2* KD are also associated with autophagy and mitophagy defects in zebrafish. Calpeptin treatment, a known activator of autophagy, is sufficient to rescue the locomotor properties, thanks to its beneficial effect on autophagy functioning in zebrafish and independently to its effect on calpain activity. LC3-II levels of primary muscle cells of TANGO2 patients are also improved by calpeptin treatment. Overall, we demonstrate that TANGO2 plays an important role in autophagy, and that autophagy efficiency is critical to prevent RM, thus giving rise to new therapeutic perspectives in the prevention of these life-threatening episodes in TANGO2 pathology.

## Introduction

Transport and Golgi Organization Protein 2 Homolog (TANGO2)-related disease is a pathology with a poor prognosis due to the recurrence of severe life-threatening rhabdomyolysis (RM) bouts ^1, 2^, characterized by the acute breakdown of skeletal myofibers ^3, 4^. Other symptoms of TANGO2 disease are developmental regression, hypoglycemia, hypothyroidism and cardiac arrhythmia, including prolonged QTc interval or ventricular fibrillation ^1, 2, 5–12^. RM can be triggered by fasting and infections, but also by exposure to cold or heat. TANGO2 function is poorly understood but has been reported to be required for endoplasmic reticulum (ER) - Golgi ^1, 12–15^, and haem ^16^ transports. Depletion of TANGO2 results in slowed cargo movements between the ER and the Golgi in patient’s fibroblasts ^14^, ER and Golgi fusion in Drosophila cells ^13^, and/or abnormal ER/Golgi morphology in different models ^1, 12, 14–16^. However, abnormal ER morphology in patients’ cells has not been systematically observed ^1, 5^. On the other hand, besides the vicinity of the ER ^12, 15^, TANGO2 is mainly cytosolic ^7^ where it has been shown to partially localize to mitochondria ^2, 9, 14, 17, 18^ and lipid droplets ^19^. Importantly, the overall abundance of major membrane and cellular lipids synthesized through ER/sarcoplasmic reticulum is significantly decreased in *tango2* mutant zebrafish at the basal level in the absence of any external trigger ^15^, and in Hep2g cells ^19^. Plasma and fibroblasts from TANGO2 patients exhibit abnormal accumulation of fatty acids and/or a defect in palmitate-dependent oxygen consumption suggesting impairment in mitochondrial fatty acid oxidation ^2, 9, 12, 17^. However, we previously showed that the main mitochondrial functions, including respiratory chain, fatty acid beta-oxidation, and Krebs cycle were normal in primary myoblasts from TANGO2 patients, implying that mitochondrial defects could be a secondary effect, with the pathogenic RM trigger remaining unknown ^5^. In a therapeutic perspective, it has recently been reported that panthotenic acid (Vitamin B5) can rescue the defects of ER to Golgi transport in TANGO2 knockout (KO) fibroblasts ^13^, as well as several behavioural traits of *tango2* knockdown (KD) Drosophila ^13^ and patients ^11, 20–22^.

Interestingly, TANGO2 patients muscle metabolism is usually sufficient to meet cellular demands apart from stress conditions. Starvation being a well-known activator of autophagy, we hypothesized that autophagy insufficiency is a critical event in RM triggering that needs to be targeted and restored to prevent these life-threatening episodes. To address this hypothesis, we examined autophagy flux in TANGO2 patients’ cells and report a defect in Microtubule-associated protein 1A/1B light chain 3B (hereafter referred to as LC3) levels upon starvation, in primary muscle cells from patients. We translated our observation from patients’ cells to *tango2* knockdown (KD) zebrafish that reproduces motor defects under basal conditions, as recently described ^15, 16^, or through addition of atorvastatin, a compound well known to induce RM. We found that *tango2* KD zebrafish display abnormal autophagy and mitophagy. Chemical activation of autophagy rescues the associated locomotor phenotype *in vivo*. In particular, calpeptin treatment rescues locomotor RM phenotype in zebrafish thanks to its beneficial effect on autophagy and independently of its effect on calpain 1/2 activity. Furthermore, calpeptin treatment also improves LC3-II levels in TANGO2 patients primary myoblasts, thus raising new therapeutic perspectives for TANGO2-related RM.

## Results

### TANGO2 plays a role at the initiation of autophagy in primary myoblasts

To determine whether TANGO2 plays a role in autophagy, we incubated primary myoblasts either in growth medium (GM) or in starvation medium (EBSS) for various durations, in the presence or absence of bafilomycin A, an inhibitor of autophagosome and lysosome fusion. Remarkably, we found a significant reduction of LC3-II absolute protein levels upon starvation in cells of three patients relative to three controls in at least three technical replicates per patient (**Fig. 1A and B**). However, p62 levels were found unchanged in TANGO2 patient myoblasts upon starvation (**Fig. S1 A and B**). Then, basal autophagy was measured by quantification of LC3-II levels over time in the presence of bafilomycin A in TANGO2 patient myoblasts and showed interpatient variability condition, as opposed to starvation-induced autophagy (**Fig. S1C and D**). Despite a downward trend, no significant difference in p62 levels was found in basal condition (**Fig. S1C and E**). Furthermore, *MAP1LC3B* (LC3B) and *SQSTM1* (p62) expressions were analyzed at the transcription level by RT-qPCR and we observed that *MAP1LC3B* expression was significantly decreased upon starvation. *SQSTM1* expression was not significantly modified in TANGO2 patients primary myoblasts, despite a decreasing tendency upon starvation. To further validate these results, LC3-II protein levels were measured upon siRNA-mediated *TANGO2* KD in control myoblasts submitted to starvation and were found decreased (**Fig. 1D-F**). Furthermore, we found that LC3-II levels were normal in TANGO2 starved fibroblasts, whereas these parameters were significantly reduced in myoblasts from the same patients, indicating potential tissue specificity for the requirement of TANGO2 in the initiation of autophagy (**Fig. S1F**).

**Figure 1.**
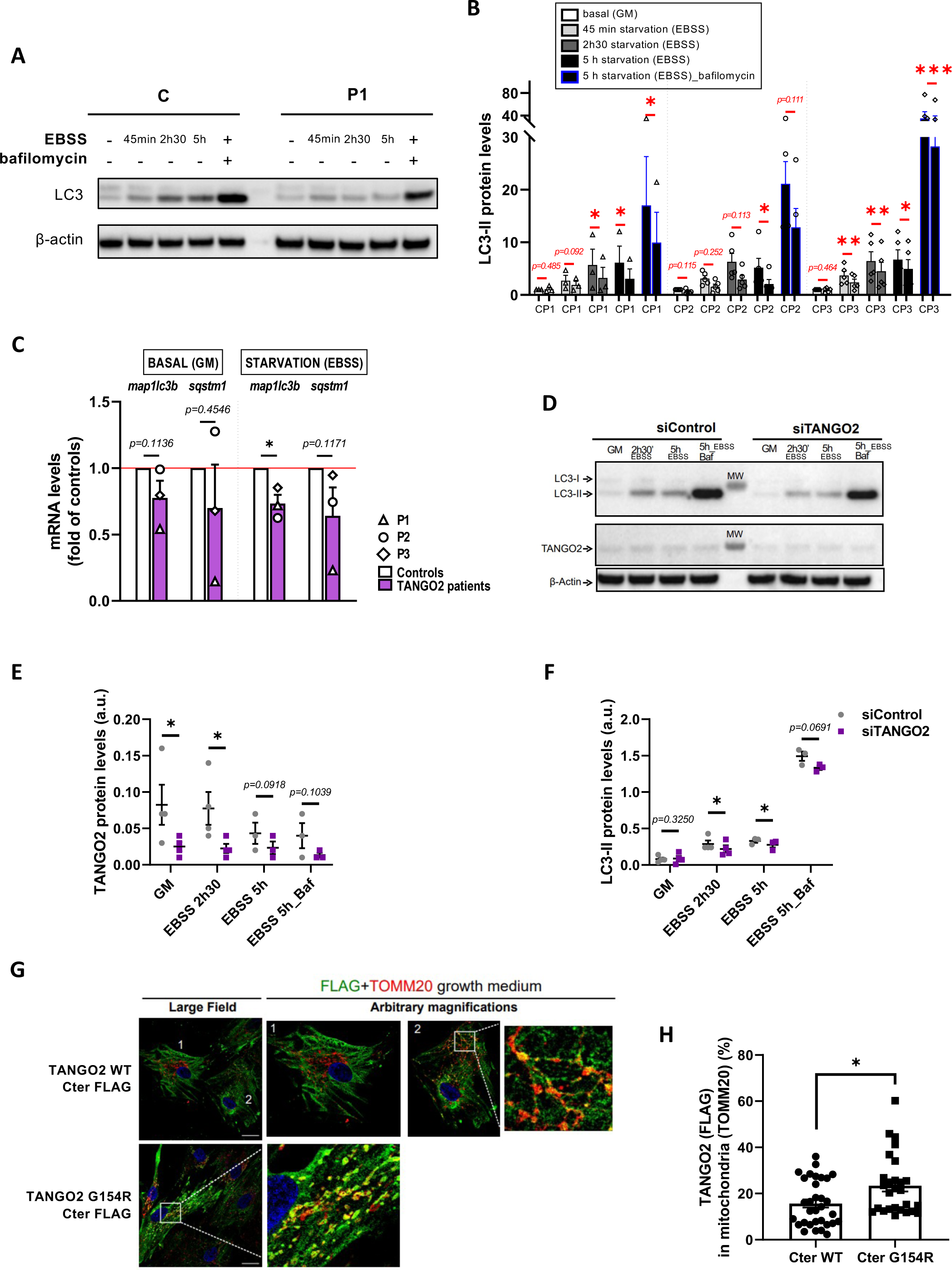
Autophagy is defective in *TANGO2* patients primary myoblasts. **A, B** Controls (C) and patients’ (P1, P2 and P3) myoblasts were cultured in Growth Medium (GM) or starved in EBSS for the indicated time points to induce autophagy. **A** Representative immunoblot showing LC3-II reduction in TANGO2 patients as illustrated in P1. **B** Quantification of LC3-II at each time point upon starvation in three differents TANGO2 patients (P1, P2 and P3). Data are shown relative to beta actin and paired between control and patient on each individual time point. n=3 patients, with at least three technical replicates per patient. *p<0.05, **p<0.01, ***p<0.001 from paired Student t-tests. **C** Expression of *MAP1LC3B* and *SQSTM1* in basal condition (GM) or upon starvation (EBSS) in patient’s primary myoblasts relative to beta actin and to the controls. Data are pooled per controls and patients and presented as mean + SEM. n=3 controls, n=3 patients with at least two technical replicates per individual. *p<0.05 or enumerated p-values from paired Student t-tests. **D-F** Control myoblasts were transfected with siRNA oligonucleotide control or against tango2, and four days after transfection, cells were cultured in Growth Medium (GM) or starved in EBSS for the indicated time points. N=4 **D** Representative immunoblots out of four independent experiments with similar results. **E**, **F** TANGO2 (**E**) and LC3-II (**F**) protein levels relative to beta actin in siControl and siTANGO2 cells. *p<0.05, or enumerated p-values from paired Student t-tests at each time point. **G, H** Patients myoblasts’ were transduced with the indicated viruses and stained against FLAG (TANGO2, green) and TOMM20 (mitochondria, red). Scale bar= 20 µm. **G** Representative images of FLAG and TOMM20 immunostainings. **H** Quantification of TANGO2 (FLAG) localization to mitochondria (TOMM20) by analysis of Mander’s coefficient of colocalization. *p<0.05 from unpaired Student t-tests.

Since the absence of TANGO2 leads to defective autophagy, we asked whether TANGO2 WT colocalizes with LC3-II autophagosomes. Using lentiviral expression of FLAG-tagged plasmids expressing WT TANGO2 or a mutant reported in patients^1^ (Glycine154Arginine, G154R) and immunolabelling, we failed to detect TANGO2 to the autophagosomes upon starvation (**Fig. S1G)**. Instead, we found that the vast majority of TANGO2 resides in the cytosol (**Fig. 1G and Fig. S1G**), in agreement with previous observations ^7^, and co-localizes little with the mitochondrial marker TOM20 (**Fig. 1G**, cell 1 **and Fig. 1H**) in partial agreement with previous data showing a mitochondrial localization and function of TANGO2 ^1, 17^. Since TANGO2 has been reported by others to play a role in mitochondria ^2, 9, 14, 17^, we further examined transduced cells to detect any partial localization of TANGO2 to mitochondria. Indeed, in a small subset of cells, TANGO2 WT partially localized to mitochondria (**Fig. 1G**, cell 2 **and Fig. 1H**). Moreover, the TANGO2 G2R Cter-FLAG mutant is present at mitochondrial vesicles (**Fig. 1G and H**). In contrast, the two TANGO2 variants with an N-terminal domain FLAG tag showed a cytosolic localization and were mostly excluded from mitochondria, irrespective of the G2R mutation (**Fig. S1H**). These data indicate that the reported mitochondrial localization of TANGO2 requires its N-terminal domain as well as unperturbed Glycine 154. Examining the expression of our constructs by Western Blotting, we noticed the presence of a band between 100-130 kDa that seems to be present only in the WT – Cter version of TANGO2 (**Fig. S1I and J**). Both FLAG and TANGO2 Proteintech antibodies can detect this band, which is not the case for lower molecular weight bands. These data suggest that TANGO2 can exist as an oligomer resistant to SDS and reducing agent, depending on the availability of the N-terminal domain of the protein.

In agreement with the minor TANGO2 mitochondrial localization at the steady state, and our previous report on normal beta-oxidation flux in patient’s myoblasts ^5^, we did not find any abnormalities of beta-oxidation flux ^23^ in whole blood of three TANGO2 patients compared to controls (**Fig. S1K**).

Taken together, these data suggest that compromised autophagy mediated by TANGO2 in muscle cells plays a pathogenic role, while in our hands, the mitochondrial function of TANGO2 described mostly in patient fibroblasts ^9, 17^ remains to be determined in muscle and blood cells.

### TANGO2 morphant zebrafish displays a RM-like locomotor phenotype due to autophagy defects

Zebrafish is a vertebrate organism widely used to model genetic conditions and major health disorders, including in the context of muscular ^24–26^ and autophagy-related diseases ^27–29^. Zebrafish skeletal muscles spontaneously start to contract as soon as 17 hours post-fertilization (hpf). By 24 hpf, myotomes are present thus enabling the embryo to coil and even respond to touch. By 48 hpf, the muscle is fully differentiated and zebrafish larvae display stereotyped escape responses to touch, allowing for an assessment of muscle performance and function using the Touch-Evoked Escape Reponse (TEER) test. We confirmed by RT-qPCR that a *TANGO2* orthologue is expressed in zebrafish at early development stages, at 1 and 2 days post-fertilization (dpf) (**Fig. 2A**). We designed two different antisense oligonucleotides coupled to the morpholino moiety (MOs) to down-regulate *tango2* expression in zebrafish: tango2-MO^atg^ is predicted to sterically prevent tango2 translation by targeting its initial AUG codon, while tango2-MO^spE3^ is predicted to alter tango2 splicing by targeting exon3-intron3 junction. Noteworthy, this splicing induction resembles the large exon3-exon 9 deletion in humans. We did not observe any developmental deficit or non-specific toxicity in the different experimental conditions as illustrated by representative zebrafish larvae of different conditions at 50 hours post fertilization (hpf) (**Fig. 2B**). In zebrafish, the RM phenotype is defined by reduced locomotion parameters and/or disrupted muscle morphology ^16, 30, 31^. Here, RM phenotype as evoked motor response was assessed through the TEER test to assess muscle performance, as previously described ^32^. Individual swimming episodes were traced for zebrafish from control-MO, tango2-MO^atg^ and tango2-MO^spE3^ (**Fig. 2C**). We observed a decreased locomotion in tango2-MO^spE3^ condition, but not in tango2-MO^atg^ condition, as compared to controls (**Fig. 2C and D**). Quantitative analysis of the TEER demonstrated that tango2-MO^spE3^ induces a locomotor phenotype, as shown by significantly decreased distance (**Fig. 2D**), which is consistent with recent observations in CRISPR KO *tango2* models ^15, 16^. To validate the specificity of this locomotor phenotype, we exposed 30 hpf zebrafish to atorvastatin (ATV, Lipitor), usually employed for the treatment of hypercholesterolemia, and proceeded to TEER analysis at 50 hpf. Statin treatments are known to induce RM, in particular in the context of an underlying genetic predisposition ^33^, and ATV has previously been shown to induce RM in zebrafish ^31^. TEER quantification demonstrated that ATV treatment exacerbates the locomotor defects of tango2-MO^spE3^ zebrafish in a dose-dependent manner (**Fig. 2E**). Indeed, while at the highest dose of 1 μM ATV, zebrafish displayed decreased swimming properties in all conditions (**Fig. 2E**), incubation with 0.5 μM ATV led to motor deficits only upon *tango2* KD, in both tango2-MO^spE3^ and tango2-MO^atg^, suggesting that another RM-specific stress is necessary to unveil RM phenotype in tango2-MO^atg^ condition (**Fig. 2E**). We took advantage of the fact that tango2-MO^spE3^ is predicted to affect tango2 expression at the mRNA level, to validate *tango2* knockdown in this condition, at 1 and 2 dpf, by gel electrophoresis of the amplicon of the targeted region (**Fig. 2E**) and by RT-qPCR (**Fig. 2F**). In addition, although tango2-MO^atg^ is predicted to inhibit *tango2* expression by sterically preventing its mRNA translation, regarding the normal locomotor behavior of tango2-MO^atg^ larvae (**Fig. 2 C, D**) and the effects of ATV treatment on this condition (**Fig. 2 E**), we looked for an eventual compensatory expression of *tango2* in tango2-MO^atg^ untreated larvae but we did not observed any difference with the controls (**Fig. 2G**), suggesting another protective mechanism in this condition.

**Figure 2.**
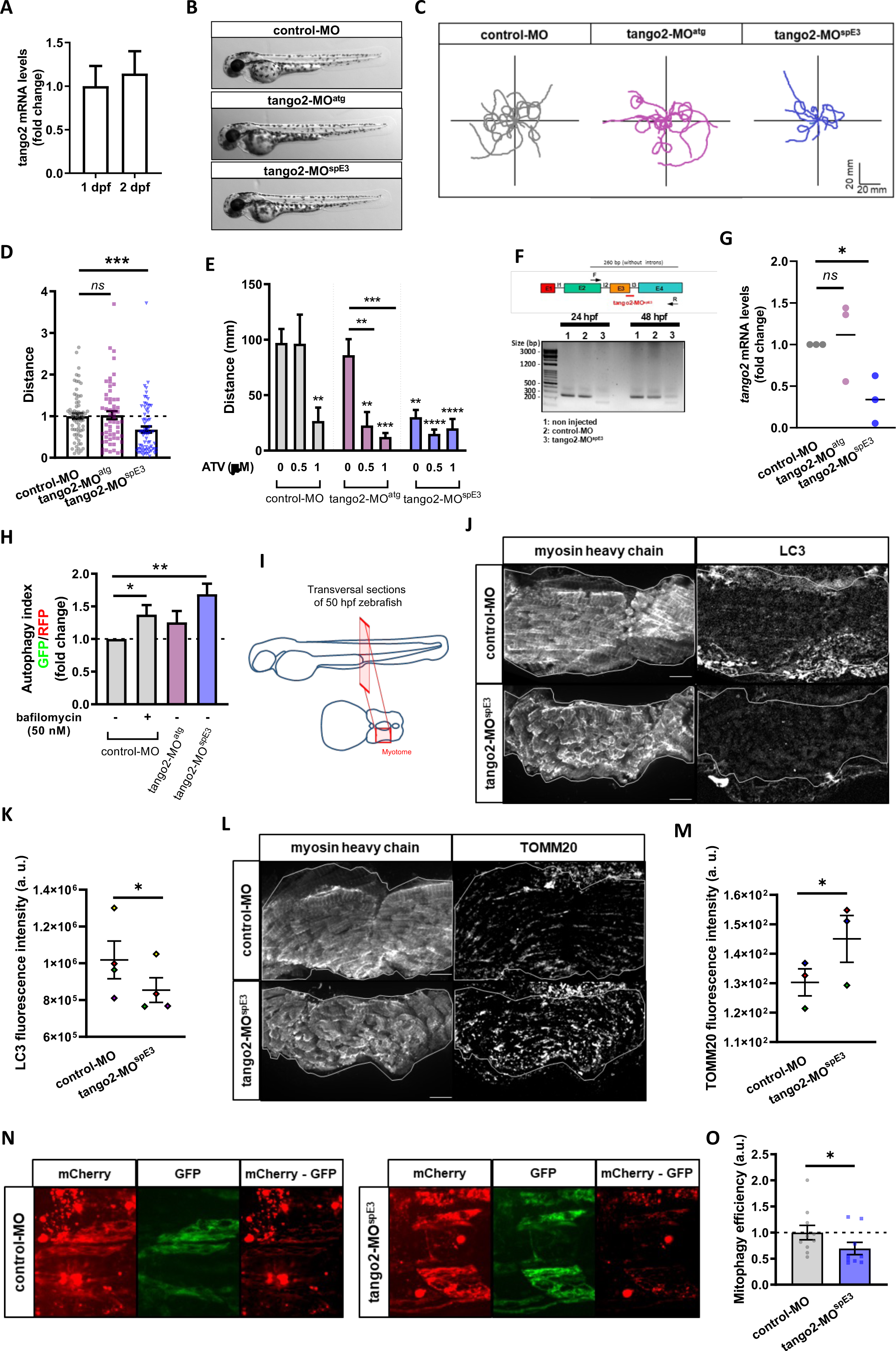
*tango2* knockdown induces a RM-like phenotype in zebrafish due to autophagy defects. **A** Expression of *tango2* zebrafish orthologue at 1 and 2 dpf embryogenic stages in zebrafish as quantified by RT-qPCR. Data are shown as mean + SEM. **B** Representative pictures of 50 hpf whole zebrafish showing no defect on global morphology. **C**,**D** TEER results**. (C)** Representative swimming trajectories of zebrafish individuals and **(D)** quantification of average swimming distance, reflecting impaired locomotor abilities of 50 hpf zebrafish from tango2-MO^spE3^ condition. Each dot represents one embryo. Data are shown as mean +/- SEM. ***p<0,001 and *ns* non-significant from Kruskal-Wallis and Dunn’s multiple comparisons test. N=5. **E** Effects of 0,5 and 1 μM atorvastatin (ATV) treatments on zebrafish swimming properties as quantified by average swimming distance parameter by TEER test. Data are shown as mean + SEM. **p<0,01; ***p<0,001; ****p<0,0001 from 2-way ANOVA and Tukey’s multiple comparison test. N=3. **F** PCR strategy and representative gel electrophoresis of amplification products from 24 and 50 hpf zebrafish showing aberrant splicing products in tango2-MO^spE3^ zebrafish at both stages. **G** tango2 mRNA levels relative to beta actin and to controls. p<0,05 and *ns* non-significant from one way ANOVA and Fisher’s post hoc analysis, N=3. **H** Monitoring of autophagy flux using the GFP-LC3-RFP-LC3Δ fluorescent probe. Autophagy index as quantified by GFP/RFP ratio by flow cytometry analysis of 30 hpf dissociated zebrafish, showing reduced autophagy flux (increased autophagy index) in tango2-MO^spE3^ zebrafish cells. Dissociated cells from control-MO zebrafish were treated with 50 nM bafilomycin as a positive control of decreased degradative efficiency of GFP-positive autophagosomes. Data are shown as mean + SEM. **p<0,01 from Kruskal-Wallis and Dunn’s test. N=8. **I** Schema of transversal sections of 50 hpf zebrafish larvae for immunostainings of the myotomes labelled with myosin heavy chain antibody. **J**, **K** Representative images (**J**) and quantification of average fluorescence intensity (**K**) from LC3 immunolabeling in myotome sections of 50 hpf zebrafish. Scale bar: 10 µm. *p<0,05 from paired Student t-test. Each pair of data is represented in the same color. **K**, **L** Representative images (**K**) and quantification of average fluorescence intensity (**L**) from TOMM20 mitochondria immunolabeling in myotome sections of 50 hpf zebrafish. Scale bar: 10 µm. *p<0,05 from paired Student t-test. Each pair of data is represented in the same color. **N**, **O** Quantification of mitophagy efficiency in zebrafish 30 hpf larvae. **N** Representative images of GFP and mCherry signals from the mitoQC probe and of the result of mCherry signal after corresponding GFP signal substraction (“mCherry - GFP”). **O** Quantification of mitophagy efficiency. p<0,05 from unpaired Student t-test (n=9).

We wondered whether autophagy disruption correlates with the locomotor phenotypes in tango2 zebrafish morphants. To test this hypothesis, we used the GFP-LC3-RFP-LC3ΔG fluorescent probe ^28^. Upon autophagy induction, the probe is cleaved, giving rise to ectopic expression of GFP-LC3 and RFP-tagged LC3ΔG mutant. GFP-LC3 is then integrated into the membrane of the autophagosome before degradation in the lysosome, whereas RFP-LC3ΔG remains in the cytosol. Thus, the autophagy index, as measured by GFP/RFP fluorescence ratio for 30 hpf zebrafish dissociated cells, inversely correlates with autophagy efficiency ^28^. We observed that the autophagy index was significantly increased in tango2-MO^spE3^ zebrafish cells, when compared to controls, indicating compromised autophagy in this condition (**Fig. 2H and S2A**). However, dissociated cells from tango2-MO^atg^ zebrafish showed no difference relative to controls, confirming that zebrafish locomotor properties correlate with autophagy efficiency. Then, we looked at endogenous levels of LC3 proteins by immunostaining of 50 hpf zebrafish transversal sections (**Fig. 2I**). Interestingly, we observed decreased LC3 levels in skeletal muscles of tango2-MO^spE3^ zebrafish, suggesting that tango2 has a role at different steps of autophagy completion (**Fig. 2 J and K**). To determine the functional implications of autophagy defects on mitochondria turnover, we immunostained transversal sections of 50 hpf zebrafish larvae with a TOMM20 antibody and we observed increased TOMM20 levels in skeletal muscles of tango2-MO^spE3^ zebrafish larvae (**Fig. 2L and M**). In order to monitor mitophagy flux in zebrafish, we used another binary-based fluorescent sensor, the ‘mito-QC’ probe ^34, 35^(**Fig 2N and O**). Mito-QC consists of a tandem mCherry-GFP tag fused to the mitochondrial targeting sequence of FIS1, a resident protein of the outer mitochondrial membrane. During mitophagy, the acidic environment of the lysosome quenches GFP fluorescence without influencing the mCherry signal. Intensity of mCherry remaining signal was measured after the substraction of the corresponding GFP signal (mCherry - GFP) and served as an indicator of mitophagy efficiency (**Fig. 2 N**). We observed that mitophagy is activated in both control and *tango2* KD conditions as indicated by the mCherry signal, but tango2-MO^spE3^ zebrafish embryos display a significantly lower mitophagy index as indicated by the decreased mCherry - GFP signal when compared to the control-MO condition (**Fig 2N and O**), thus confirming that the mitophagy process is impaired upon *tango2* KD.

### Calpeptin treatment rescues TANGO2 pathology in vivo and in vitro

To counteract autophagy defects in tango2-MO^spE3^ zebrafish, we treated 30 hpf larvae with calpeptin, a calpain inhibitor known to activate autophagy including in zebrafish ^36^ and previously used as a potential therapeutic in a Machado-Joseph zebrafish model ^37^. Calpeptin treatment was sufficient to normalize locomotor parameters of tango2-MO^spE3^ zebrafish, and increased the travelled distance in treated zebrafish significantly (**Fig. 3A**). Importantly, we observed that the autophagy flux of tango2-MO^spE3^ cells is also improved by calpeptin treatment (**Fig. 3B**). Calpains are a family of calcium-dependent proteases having multiple important cellular roles, including autophagy regulation ^38, 39^. In particular, calpains are known to induce the cleavage of ATG5 (33 kDa full length), giving rise to a truncated 24 kDa form that can bind to mitochondria and promote apoptosis ^38, 39^. Then we investigated whether calpains 1 and 2 activities were modified in our zebrafish model using an enzyme activity fluorescent assay. While we observed that calpains activities were significantly decreased by calpeptin treatment, we did not observe any difference between untreated control and *tango2* KD conditions (**Fig. 3C**). Furthermore, cleavage of ATG5 (33 kDa) to 24 kDa form was measured by Western Blot of zebrafish proteins (**Fig 3C and D**). Consistent with calpains 1/2 activity results, calpeptin treatment reduced the cleavage of ATG5 as demonstrated by the significantly decreased ratio of 24/33 kDa forms levels but we did not observe any difference between control-MO and tango2-MO^spE3^ conditions on ATG5 cleavage (**Fig. 3C and D**). Overall, this suggests that the beneficial effect of calpeptin treatment on autophagy functioning and locomotor phenotype in *tango2* KD condition is independent of its effect on calpain activity. Then, we treated zebrafish larvae with two other activators of autophagy, rapamycin (**Fig. S2B**) and torin1 (**Fig. S2C**) and we observed that both drugs were able to ameliorate the travelled distance of tango2-MO^spE3^, thus reinforcing the notion that autophagy activation is responsible for the rescue of the RM phenotype associated with *tango2* KD.

**Figure 3.**
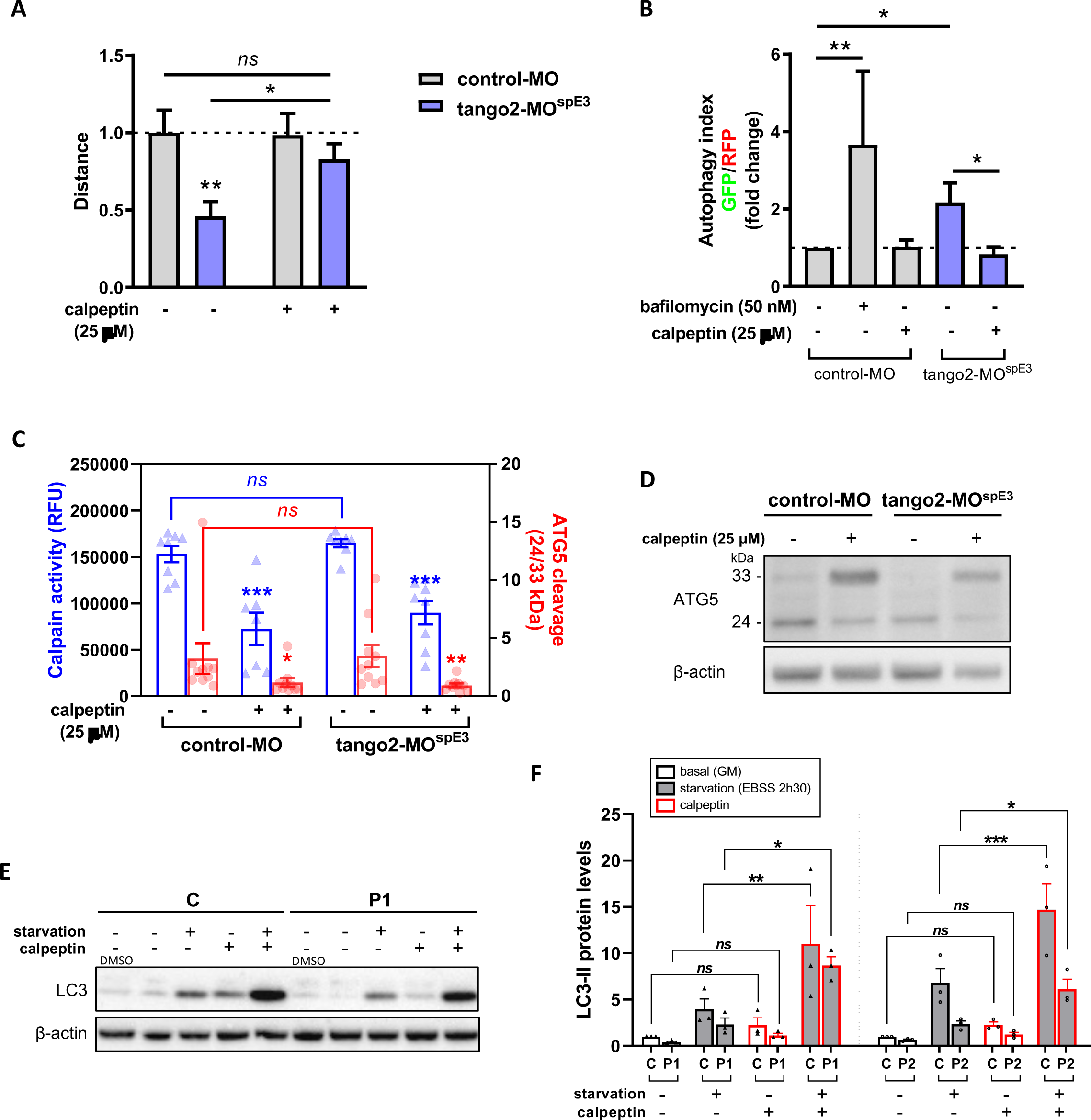
Calpeptin ameliorates TANGO2 disease features in zebrafish and in primary myoblasts. **A.** Average swimming distance from TEER test showing the beneficial effect of 25 μM calpeptin treatment on RM-like features of 50 hpf zebrafish. Data are shown as mean + SEM. *p<0,05; **p<0,01 and *ns* non-significant from 2-way ANOVA and Tukey’s test. N=4. **B.** Autophagy index of 30 hpf zebrafish dissociated cells treated with 25 μM calpeptin, showing that calpeptin restores autophagy efficiency of tango2-MO^spE3^ zebrafish. 50 nM bafilomycin was used as a positive control of decreased degradative efficiency of GFP-positive autophagosomes. Data are shown as mean + SEM. *p<0,05; **p<0,01 from 2-way ANOVA and Tukey’s test. N=6. **C** Measure of calpains 1/2 activity in 50 hpf zebrafish proteins by fluorescent enzymatic assay (plotted in blue on the left Y axis) and by Western Blot analysis of ATG5 cleavage (plotted in red on the right Y axis). *ns* non significant and *p<0,05; **p<0,01 ***p<0,001 from 2-way ANOVA followed by Tukey’s multiple comparison analysis for both datasets. **D** Representative ATG5 immunoblot from a single experiment out of 10 independent experiments with similar results. **E**, **F** Western Blot results of control and TANGO2 patients primary myoblasts cultured in growth medium (GM) or starved in EBSS, treated or not with 50 μM calpeptin. **E** Representative LC3 immunoblot from a single experiment out of 9 independent experiments with similar results. **F** LC3-II protein levels showing that calpeptin treatment improves LC3-II levels upon starvation in control and TANGO2 primary myoblasts. Calpeptin treatment effect was analyzed on GM and starvation conditions. *p<0.05 and *ns* non-significant, from 2-way ANOVA and Fisher’s post-hoc analysis.

Having discovered that inducing autophagic in zebrafish by calpeptin treatment reverts the disease phenotype in *tango2* deficient zebrafish, we tested whether calpeptin treatment can improve the defective LC3-II levels in primary TANGO2 patient myoblasts (**Fig. 3E and F, Fig. S2D**). Calpeptin restored LC3-II levels in primary TANGO2 KO myoblasts submitted to starvation (**Fig. 3E and F, Fig. S2D**), demonstrating that calpeptin treatment can improve autophagy functioning in human cells.

## Discussion

Here, we report that TANGO2 protein has a role in the regulation of autophagy. Indeed, we observed that TANGO2 deficiency results in reduced LC3-II levels upon starvation in primary muscle cells from patients, and in defective autophagy in zebrafish. Indeed, TANGO2 patients exhibit normal or subnormal muscle function between RM episodes that are worsened by extrinsic triggers such as fasting, a well-known activator of autophagy. Functional autophagy is essential for the maintenance and adaptation to stress of muscle cells, and defects in all steps of this pathway have been associated with skeletal muscle diseases ^40–45^, including metabolic RM ^46, 47^. Supporting that, *tango2* KD led to reduced evoked swimming parameters in zebrafish, which is consistent with recent studies showing a myopathy-like phenotype in zebrafish upon *tango2* knockout ^15, 16^. We also show that incubation of *tango2* KD zebrafish with atorvastatin, a drug known to induce RM ^33, 48^ including in zebrafish ^31, 49^, exacerbates the motor deficits observed in *tango2* KD, thus confirming the specificity of *tango2* KD phenotype. Furthermore, we corroborate *in vivo* that the RM-like phenotype induced by *tango2* loss of function correlates with impaired autophagy function, as demonstrated by the decreased degradation of autophagosomes containing exogenous LC3 and by the decreased levels of LC3 proteins in *tango2* KD zebrafish. Our observation that the starvation-induced autophagy does not show any alteration in skeletal fibroblasts from the same TANGO2 deficient patients indicates that the RM pleiotropy of TANGO2 pathology could be due to tissue-specificity of autophagy dysfunction in organs at risk of decompensation during fasting.

Importantly, we demonstrated that restoring autophagy function by calpeptin, a calpain inhibitor, is sufficient to rescue the RM-like phenotype in *tango2* KD zebrafish in a calpain-independent way, as well as to improve LC3-II levels in starved TANGO2 primary myoblasts. Importantly, the improvement of locomotor function of *tango2* KD zebrafish by two other autophagy activators, rapamycin and torin1, confirms that autophagy impairment is an important mechanism in TANGO2 pathophysiology

There is a functional link between the COPII machinery involved in anterograde trafficking, that is known to be impaired in TANGO2 disease, and autophagy, as COPII vesicles serve as a membrane precursor for autophagosome biogenesis ^50–55^. TRAPPopathies due to *TRAPPC11* or *TRAPPC12* mutations, that present neurological features associated with possible rhabdomyolysis, display a defect in both trafficking machinery between the ER and the Golgi apparatus and autophagy ^56–58^, similarly to TANGO2 disease. On the other hand, recent advances in the understanding of TANGO2 disease point towards defects in lipid metabolism with decreased overall abundances of membrane and cellular lipids ^13, 15, 16, 19^. Indeed, two recent papers have shown abnormal lipid profiles in TANGO2 models, with reduced levels of phosphatidic acid and increased levels of its unacetylated precursor, lysophosphatidic acid, in Hep2g cells ^19^, and decreased phosphatidylcholine and triglycerides in zebrafish ^15^. In this sense, we and others have previously demonstrated that mutations in the phosphatidate phosphatase *LPIN1*, another genetic cause of RM predisposition, lead to a selective loss of phosphatidylinositol-3-phosphate ^46, 59^, which is particularly enriched in autophagosomes and involved in all steps of autophagy ^60^, with subsequent autophagy impairment ^46, 59^. Abnormal phospholipids and autophagy defect have been also associated with other skeletal myopathies such as Vici syndrome ^47^. Overall, this raises the hypothesis that mutations in genes involved in lipid metabolism can converge to autophagy insufficiency which can be exacerbated by stress conditions, such as starvation, thus leading to RM outbreak. Altogether, considering that i) starvation affects lipid metabolism and autophagy, ii) membrane lipid composition is critically involved in autophagy regulation and processing ^61^, iii) the ER-ERGIC-Golgi axis provides membranes to form autophagosomes, thereby connecting vesicular forward transport with autophagy, iv) the reports on defective lipid metabolism in TANGO2 KO cells and animal models, and v) the autophagy defect in TANGO2 deficient models observed in this study, it is likely that TANGO2 is involved in the anterograde trafficking and in the metabolism of lipids of membrane structures participating in autophagy and that TANGO2-related muscle symptoms are the consequence of insufficient autophagy in patients exposed to stress condition, such as starvation triggering autophagy.

In physiological conditions, we observe that the vast majority of TANGO2 localizes to the cytosol and only a minor portion to mitochondria, in agreement with two previous reports ^14, 16^. These observations are in line with our previous study, where we did not observe any anomalies in metabolic beta-oxidation flux in such cells upon palmitate loading, or in mitochondrial structure and function ^5^, and with the results presented here showing normal mitochondria beta-oxidation flux in whole blood of TANGO2 deficient patients. Other studies also found normal OXPHOS in patient’s cells ^7^ or in TANGO2 iPSC derived cardiomyocytes ^62^. However, abnormal acylcarnitine profiles in plasma of some patients ^1, 2, 9, 12^ and ultrastructure defects in mitochondria ^15^ or abnormal mitochondrial findings ^7, 9, 13, 17, 19, 63^ have been also previously reported, in accordance with our finding of impaired mitophagy process upon *tango2* KD. These discrepencies suggest that mitochondrial dysfunction is secondary in the pathophysiology of TANGO2 disease. Accordingly, we found that overexpressed TANGO2 G154R -FLAG mutant ^1^ in myoblasts localizes more prominently to mitochondria than its WT counterpart. On the basis of these observations, we speculate that TANGO2 can be constitutively recruited to mitochondria via its mitochondrial localization signal at the N-terminus ^14^ and cycle back to the cytosol. The mutation G154R might act as an anchor to a mitochondrial protein, preventing its release to the cytosol.

Consequently, the reported restoration of CoA by vitamin B5 ^11, 13, 20–22^, the obligate precursor of CoA, an essential pathway conserved in both prokaryotes and eukaryotes, may concern cytosolic CoA, as the intramitochondrial coenzyme A pool is not limiting for the mitochondrial beta-oxidation pathway. A possible way to reconcile these observations made in different models would place TANGO2 as an important protein in the cytosol, notably as an intermediate in cytosolic CoA formation, thereby acting as an upstream membrane precursor to form COPII vesicles or autophagosomes. Interestingly, immunofluorescence experiments in primary myoblasts ruled out a direct TANGO2 localization to autophagosomes under starvation. Under nutrient-rich conditions, constitutive ER to Golgi transport is already affected in TANGO2 KO cells by the constitutive nature of this process ^14^. Conversely, basal autophagy was broadly normal but with variability from patient to patient, suggesting that the distinct severity of TANGO2 patients’ symptoms may rely on differentially affected autophagy under steady state.

Finally, recent work has reported that TANGO2-GFP is expressed at the expected size in various cell lines ^14, 19^. Our observation of a band between 100-130 kDa detectable upon prolonged exposure in control but not patient cells might indicate that TANGO2 can oligomerize, although at a very minor rate. Further biochemical studies, as well as the development of antibodies against TANGO2 that work in immunofluorescence staining would be instrumental for studying this hypothesis.

In conclusion, our results support that abnormal autophagy is a critical mechanism in TANGO2-related RM pathophysiology that needs to be targeted in a therapeutic perspective. In addition, we report several autophagy activators as candidate for prevention of RM bouts in TANGO2 patients.

## Methods

### Patient myoblasts

Myoblasts from three patients (P1, P2 and P3) were obtained from skeletal muscle biopsy from individuals (deltoid region) and 3 age-matched control individuals (paravertebral region). Human primary myoblasts were isolated and grown as described ^46^. To mimic starvation, growth medium (GM) was replaced by EBSS medium. Calpeptin (2h30 treatment) and bafilomycin A1 were applied in the medium at the final concentration of 25 µM and 100 nM, respectively. The patients harbored pathogenic variants in *TANGO2* gene (Patient 1 and Patient 3: homozygous deletion Exon 3-Exon 9, Patient 2: homozygous c.262C>T (p.Arg88) found by next generation sequencing (NGS) ^5^. Informed consent was obtained from TANGO2 patients and controls after obtaining the ethics approval to work on human samples by the *Comité pour la protection des personnes* (CPP, 2016) and the declaration of human myoblasts to the *Département de la Recherche Clinique et du Développement*.

### Plasmids, lentivirus production, and transduction

Codon optimized 2X FLAG TANGO2 WT (NP_001270035) was inserted into pMK-T or pLVX-EF1α-IRES-puro with restriction sites SpeI/NotI by GeneArt (Thermosfisher, Germany). The mutant 2X FLAG TANGO2 Gly154Arg was generated with the In-Fusion HD Cloning Plus Kit (Takara Bio Europe) following the manufacturer’s instructions using the pMK-T plasmid as template, and then cloned into pLVX-EF1α-IRES-puro, to produce pLVX 2X FLAG TANGO2 G2R. To generate C-terminal fusion proteins, the FLAG tag was removed as above, serving as a template to generate pMK-T TANGO2 WT-2X FLAG C-ter or pMK-T TANGO2 G2R-2X FLAG C-ter. Subsequently, the inserts were cloned into pLVX-EF1α-IRES-puro to produce pLVX TANGO2 WT-2X FLAG Cter or pLVX TANGO2 G2R-2X FLAG Cter. Plasmid DNA was extracted and purified with Nucleobond Xtra Maxi EF kit (Macherey-Nagel) and used to produce lentiviruses with an average titer of 10^9^ TU/ml. Primary myoblasts at early passages were transduced at a MOI=50 in the presence of polybrene at 8 μg/ml, overnight. Next day, the medium was changed, and two days later, puromycin was added at a final concentration of 5 μg/ml. Medium was changed routinely in the presence of puromycin, and the cells were passaged at least twice before an experiment.

### siRNA knockdown in myoblasts

Control myoblasts were seeded into 6-well plates and transfected with 100 nM control siRNA D-001810-10-05 (Horizon) *or* siRNA against Tango2 *L-* 016397-02-0005 (Horizon), with Dharmafect transfection reagent (T-2001-01 Horizon, 5 uL/well).

### Gene expression analysis in myoblasts

Total RNA was isolated from skeletal muscle using NucleoSpin RNAXS kit (Macherey-Nagel). Single-stranded cDNA was synthesized from 1 μg of total RNA using the High Capacity RNA-to-cDNA Kit (Applied biosystems) after depleting genomic DNA. The expression of *MAP1LC3B* and SQSTM1 genes in primary myoblasts was assessed by RT-qPCR using Power SYBR® Green PCR Master Mix and normalized against β*-actin*. Primers are the followings: Reactions were performed in triplicate on an Azure Cielo Real-Time PCR machine (Azure Biosystem). The RQ value was equal to 2ΔΔct where ΔΔct is calculated by (Ct target-Ct β-actin) test sample - (Ct target-Ct β-actin) calibrator sample. Each value was derived from three technical replicates.

### Western Blot (cells)

Myoblasts or fibroblasts were lysed in RIPA buffer with protease inhibitors for 10 minutes on ice, prior to sonication (5 pulses, 5 seconds), and further 10 minutes on ice. Post-nuclear supernatants were quantified by BCA before loading equal amounts of protein per lane onto NuPage 4-20% Bis-Tris gels (ThermoScientific). Dry transfer onto PVDF membranes was done with the iBlot2 device according to the manufacturer’s instructions (ThermoScientific). Primary antibodies anti-LC3B (clone 4E12, MBL International, 1:1000), anti-p62 (H00008878-M01 Abnova), anti-FLAG (Sigma 1:1000), anti-TANGO2 (Proteintech 27 846-1-AP 1:500) and anti-β-actin (sc-81178, Santa Cruz Biotechnology, 1:1000). Immunoblots were incubated with the corresponding secondary horseradish peroxidase (HRP) conjugated antibodies, signal was enhanced with ECL (enhanced chemilunescence) and detected by a ChemiDoc^TM^ Imaging system (Bio-rad).

### Confocal microscopy

Cells seeded onto glass cover slips were fixed with 2% paraformaldehyde, quenched with 300 mM glycine, and permeabilized using 0.2% saponin and 0.2% BSA in PBS. Primary antibodies were diluted in permeabilization buffer (LC3B 1:100, TOM20 (Santa-Cruz rabbit ref, FLAG 1:300), and secondary antibodies to 1:300. Nuclei were stained with DAPI (100 ng/ml). Image acquisitions were performed with a 63× oil immersion objective (NA 1.4) through a laser scanning confocal microscope (TCS SP8-3X STED; Leica Microsystems). Images were processed with FIJI 1.83 software ^64^. Colocalization analysis was performed with JACoP from ImageJ, throughout the entire volume of each cell, unless otherwise specified. Data are reported as the Mander’s coefficient.

### Zebrafish Maintenance

Adult and larval zebrafish (*Danio rerio*) were maintained at Imagine Institute (Paris) facility and bred according to the National and European Guidelines for Animal Welfare. Experiments were performed on wild type and transgenic zebrafish larvae from AB strains. Zebrafish were raised in embryo medium: 0,6 g/L aquarium salt (Instant Ocean, Blacksburg, VA) in reverse osmosis water 0,01 mg/L methylene blue. Experimental procedures were approved by the National and Institutional Ethical Committees. Zebrafish were staged in terms of hours post fertilization (hpf) based on morphological criteria and manually dechorionated using fine forceps at 24 hpf. All the experiments were conducted on morphologically normal zebrafish larvae.

### Microinjections

Morpholino antisense oligonucleotides (MO; GeneTools, Philomath, USA) were used to specifically knockdown the expression of *tango2* zebrafish orthologue. The MOs were designed to bind to the ATG (TANGO2-MO^atg^) or to a splicing region in exon3 (TANGO2-MO^spE3^). The sequences are respectively: 5’-ACTTCAAGAAGATGATGCACATGAG-3’ and 5’-ATAAGGATGATATTTACCGCTGAGG-3’. Control morpholino (control-MO), not binding anywhere in the zebrafish genome, has the following sequence 5’-CCTCTTACCTCAGTTACAATTTATA-3’. The GFP-LC3-RFP-LC3ΔG autophagy probe ^27^ was injected at a final concentration of 120 ng/μL along with the amo. The mitoQC probe ^34, 35^ was injected at a final concentration of 100 ng/uL. All the microinjections were carried out at one cell stage.

### Gene expression analysis in zebrafish

Total RNA was isolated from injected fish using TRIzol Reagent (Sigma) according to the manufacturer’s protocol. First-strand cDNAs were obtained by reverse transcription of 1 μg of total RNA using the High-Capacity cDNA Reverse Transcription Kit (Roche), according to the manufacturer’s instructions. Quantitative PCR amplification was performed with SyBer2X Gene Expression Assays using the following primers; (Fw) 5’ TCTTGAAGTTCGACCCTCGGC 3’ and (Rv) 5’ CAAAAAACCTCTCCCCTGGGC 3’. Data were analyzed by transforming raw Ct values into relative quantification data using the delta Ct method. To assess the efficiency of tango2-AMO^spE3^ on tango2 exon 3 splicing, the same primers were used (the reverse primer targets exons 4 and 5). The PCR product was loaded on a 1% agarose gel.

### Zebrafish locomotor analysis

Locomotor behavior of 50 hpf zebrafish larvae were assessed using the Touched-Evoked Escape Response (TEER) test. Briefly, zebrafish were touched on the tail with a tip and the escape response were recorded using a Grasshopper 2 camera (Point Grey Research, Canada) at 30 frames per second. Travelled distance was quantified frame per frame for each embryo using the video tracking plugin of FIJI 1.83 software ^64^. For drugs treatment experiments, 30 hpf zebrafish embryos were raised in embryo medium containing 25 μM calpeptin, 0,5 μM or 1 μM atorvastatin, 0,5 µM rapamycin and 0,15 µM torin1 dissolved in DMSO, and locomotor phenotype was assessed at 50 hpf as described above.

### Autophagy analysis in zebrafish

To monitor autophagy flux in zebrafish, we co-injected the GFP-LC3-RFP-LC3ΔG probe developed by Mizushima’s laboratory ^28^ with control-MO, tango2-MO^atg^ or tango2-MO^spE3^. Dechorionated 30 hpf zebrafish were dissociated in EDTA-trypsin 0,25% at 28°C and by trituration. Digestion was stopped with 10% fetal calf serum and suspended cells were strained with a 40 μM strainer. Cells were then centrifuged (5 min at 3000 rpm, 4°C), washed and resuspended with cold HBSS, twice. As a proof of principle, we treated dissociated cells from control-MO condition with 50 nM bafilomycin A1 for 1 hour. Autophagy was activated in dissociated cells with 25 uM calpeptin for 1 hour. The proportions of GFP-positive cells and of RFP-positive cells were quantified by flow cytometry, as previously established ^28^ using a MACSQuant® Analyzer 10 Flow cytometer (Miltenyi Biotec, Germany). Dissociated cells from 30 hpf non injected zebrafish were used as a negative control for fluorescence and compensation was made with cells from 30 hpf dissociated zebrafish expressing GFP or RFP fluorescence only. Data were processed using FlowJoTM 10 (BD, USA).

### Mitophagy analysis in zebrafish

To monitor mitophagy flux in zebrafish, we co-injected the mito-QC probe with control-MO or tango2-MO^spE3^. GFP and mCherry signals were captured from myotomes of alive 30 hpf zebrafish larvae with a Spinning Disk system (Intelligent Imaging Innovations, USA), an Examiner.Z1 upright stand (Carl Zeiss, Germany), a CSU-W1 head (Yokogawa, Japan), and an ORCA-Flash 4.0 camera (Hamamatsu, Japan), with a 20x objective (NA1). Stacks were processed with FIJI 1.47 software and same treatments were applied for all the conditions. Maximal projections images were created from Z-stacks. As an indicator of mitophagy efficiency, remaining raw mCherry fluorescence intensities were measured after substraction of the corresponding GFP signal in two randomly selected regions of interest (ROI) per embryo with at least nine embryos per condition, using FIJI 1.83 software calculator plugin^64^.

### Measure of calpains by enzymatic assay and ATG5 cleavage in zebrafish proteins

Proteins were extracted from 50 hpf zebrafish embryos using the lysis buffer from the calpain 1/2 activity kit (InnoZyme™ Calpain 1/2 Activity Assay Kit CBA054 purchased from Merck Millipore). Protein concentration was adjusted to 2 mg/mL and calpain enzymatic activity was measured in microtubes following the manufacturer instructions, using supplied positive and negative regulators of calpain activity. Fluorescent intensity was measured with a Clariostar (BMG Labtech). ATG5 cleavage was analyzed from the same protein samples by Western Blot. 25 ug of proteins were separated by SDS polyacrylamide gel electrophoresis. Samples were denatured at 98°C for 5 minutes. Separated proteins were transferred to nitrocellulose membranes and probed with anti-ATG5 (E-AB-10814-60 Quimigen) primary antibody. Anti-beta-actin (4970S Cell Signaling Technology) was used as loading control. The blots were incubated with the corresponding secondary HRP-conjugated antibodies, signal was enhanced with ECL and detected by a ChemiDoc^TM^ Imaging system (Bio-rad). Ratio of bands intensity were measured with FIJI 1.83 ^34, 35, 38, 39, 64, 65^.

### Beta oxidation flux

Beta oxidation flux was measured in whole blood by a method adapted from Dessein et al ^23^. Briefly, deuterated acylcarnitine produced after incubation of whole blood with carnitine and deuterated palmitate ([16-^2^H_3_, 15-^2^H_2_, 14-^2^H_2_, 13-^2^H_2_]-palmitate) were measured by LC-MS/MS.

### Immunofluorescent staining on zebrafish larvae transversal sections

50 hpf embryos were anaesthetized in 0,2 % tricaine, fixed in 4% PFA and prepared for cryosections as previously described ^65^. Samples were cut into 12 μm thick transversal sections which were blocked and permeabilized with 0.2% gelatin, 0.25% Triton X-100 (diluted in 1X PBS and incubated overnight with primary antibodies: anti-myosin heavy chain (1:25 DSHB MF20) to label skeletal muscles, anti-LC3 (1/200, Novus Biologicals NB100-2220), anti-TOMM20 (1:200, rabbit polyclonal, HPA011562 Sigma Aldrich, USA). After several washes, sections were incubated 1 hour with the appropriate secondary antibodies conjugated to an Alexa Fluor® (1:500, Thermo Fisher Scientific). Sections were rinsed and mounted in Fluoromount-G™ medium (Thermo Fisher Scientific). Images of myotomes area were captured with a Spinning Disk system (Andor technology, UK; Leica Microsystems, Germany), a DMI8 inverted stand (Leica Microsystems, Germany), a CSU-X head (Yokogawa, Japan) and a QE-180 camera (Hamamatsu, Japan), with a 63X objective. Stacks were processed with FIJI 1.83 software and same treatments were applied for all the conditions. Maximal projections images were created from the same number of Z-stack images per embryo. LC3 and TOMM20 average fluorescence intensities images were measured in eight random ROI in myotomes of four and three different embryos per condition, respectively.

### Statistical analysis

Data were plotted and analysed using Prism (GraphPad, USA). Statistical details are indicated in the legends.

## Abbreviations

ATV: atorvastatin
dpf: days post-fertilization
ER: endoplasmic reticulum
hpf: hours post-fertilization
KD: knockdown
KO: knockout
LC3: Microtubule-associated protein 1A/1B light chain 3B
MO: morpholino
RM: rhabdomyolysis
TANGO2: Transport and Golgi Organization Protein 2 Homolog

## Acknowledgements

The authors thank the zebrafish platform and the cell-imaging platform, for technical assistance in confocal imaging acquisition and analysis; Etienne Morel, Perrine Renard, for insightful discussions. Lentiviruses were produced by the platform Structure Fédérative de Recherche Necker Vecteurs viraux et transfert de genes (Structure Féderative de Recherche Nêcker-Université Paris Descartes, S Fabrega). This work was supported by grants from Fondation Lejeune 2017 to PdL, Agence Nationale de la Recherche to PdL and PvE (ANR – AAPG 2018 CE17 MetabInf; ANR – AAPG 2023 CE17 Rhabdophagy), the Association Française contre les Myopathies to PdL, SM, HdC and EK (AFM 2016 – 2018 n°19773; AFM 2022 – 2025 n°24269), Tango2 Research Foundation 2020 to PdL, SM, HdC and EK, Prix Necker 2017 to PdL, Prix Sauver la vie Université Paris Cité 2020 to PdL, and patient associations (No Myolyse, Des ailes pour L, Nos Anges, OPPH, de Miniac en attente, Hyperinsulinisme, Noa Luû). SM was supported by a funding from the ANR – AAPG 2018 CE17 MetabInf. HdC is supported by a funding from the ANR – AAPG 2023 CE17 Rhabdophagy.

## Disclosure statement

The authors have no conflict of interest.

## Notes

### Competing Interest Statement

The authors have declared no competing interest.

### Summary of Updates

error in figures order

