## Supplementary Figures for "TANGO2-related rhabdomyolysis symptoms are associated with abnormal autophagy functioning"

### Slide 1
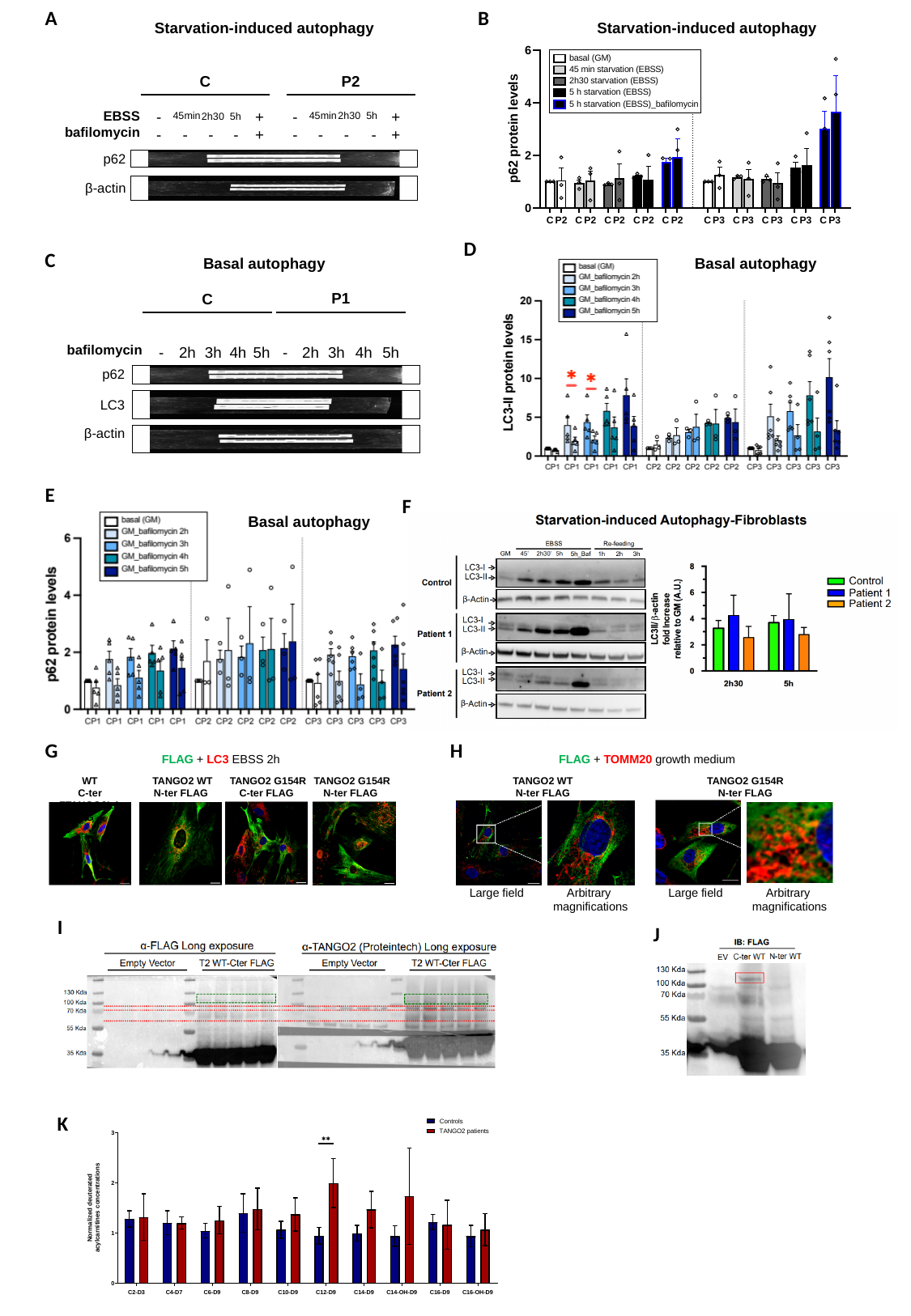

A
B
Starvation-induced autophagy
Starvation-induced autophagy
P2
C
-
-
-
-
-
+
+
-
-
-
-
-
+
+
EBSS
bafilomycin
45min
2h30
5h
45min
2h30
5h
p62
β-actin
D
C
Basal autophagy
Basal autophagy
P1
C
-
2h
3h
4h
5h
-
2h
3h
4h
5h
bafilomycin
p62
LC3
β-actin
E
F
Basal autophagy
G
H
FLAG + TOMM20 growth medium
FLAG + LC3 EBSS 2h
TANGO2 WT
N-ter FLAG
WT
C-ter FTANGO2LAG
TANGO2 WT
N-ter FLAG
TANGO2 G154R
C-ter FLAG
TANGO2 G154R
N-ter FLAG
TANGO2 G154R
N-ter FLAG
Large field
Arbitrary
magnifications
Large field
Arbitrary
magnifications
I
J
K

### Slide 2
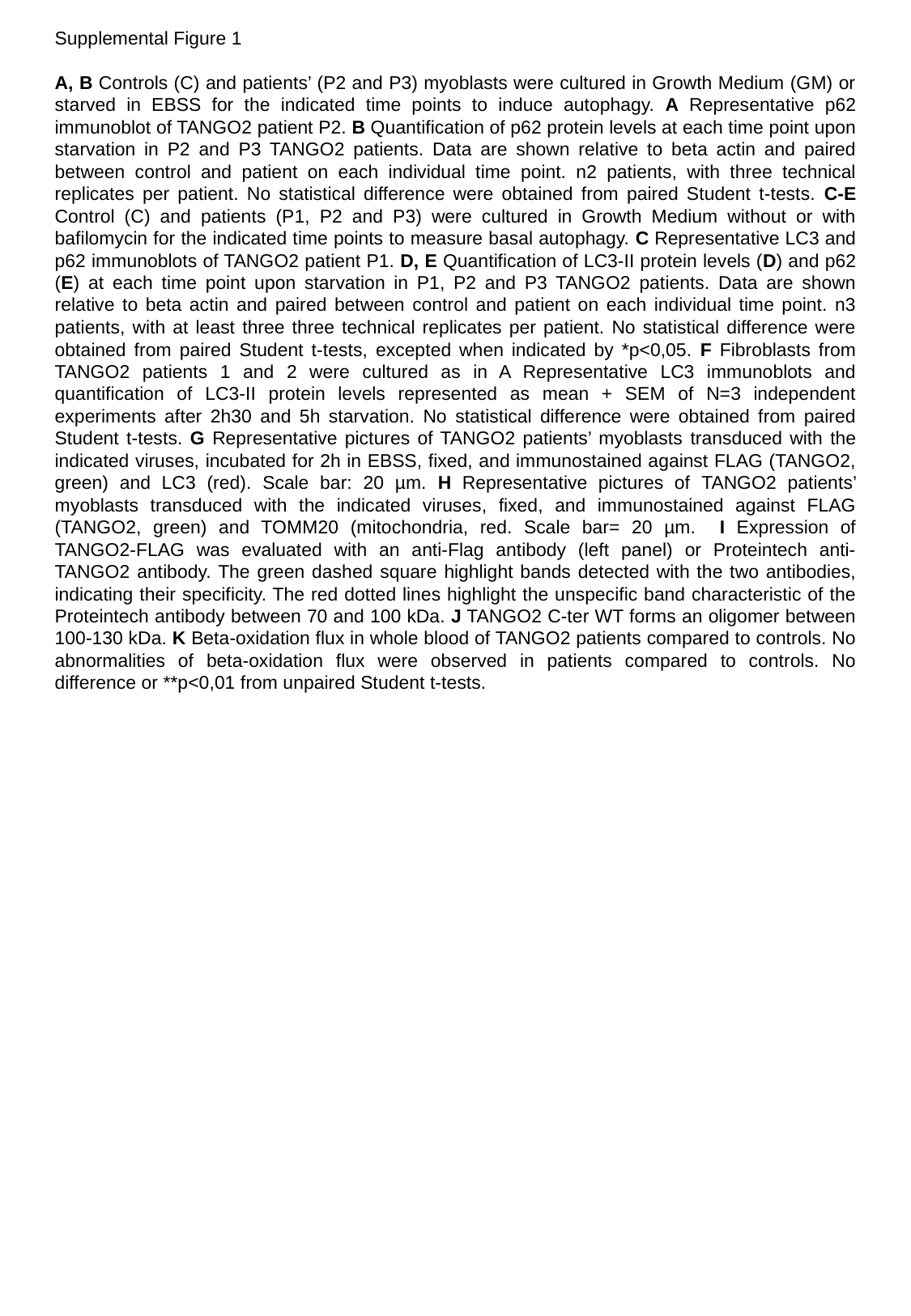

Supplemental Figure 1
A, B Controls (C) and patients’ (P2 and P3) myoblasts were cultured in Growth Medium (GM) or starved in EBSS for the indicated time points to induce autophagy. A Representative p62 immunoblot of TANGO2 patient P2. B Quantification of p62 protein levels at each time point upon starvation in P2 and P3 TANGO2 patients. Data are shown relative to beta actin and paired between control and patient on each individual time point. n2 patients, with three technical replicates per patient. No statistical difference were obtained from paired Student t-tests. C-E Control (C) and patients (P1, P2 and P3) were cultured in Growth Medium without or with bafilomycin for the indicated time points to measure basal autophagy. C Representative LC3 and p62 immunoblots of TANGO2 patient P1. D, E Quantification of LC3-II protein levels (D) and p62 (E) at each time point upon starvation in P1, P2 and P3 TANGO2 patients. Data are shown relative to beta actin and paired between control and patient on each individual time point. n3 patients, with at least three three technical replicates per patient. No statistical difference were obtained from paired Student t-tests, excepted when indicated by *p<0,05. F Fibroblasts from TANGO2 patients 1 and 2 were cultured as in A Representative LC3 immunoblots and quantification of LC3-II protein levels represented as mean + SEM of N=3 independent experiments after 2h30 and 5h starvation. No statistical difference were obtained from paired Student t-tests. G Representative pictures of TANGO2 patients’ myoblasts transduced with the indicated viruses, incubated for 2h in EBSS, fixed, and immunostained against FLAG (TANGO2, green) and LC3 (red). Scale bar: 20 µm. H Representative pictures of TANGO2 patients’ myoblasts transduced with the indicated viruses, fixed, and immunostained against FLAG (TANGO2, green) and TOMM20 (mitochondria, red. Scale bar= 20 µm. I Expression of TANGO2-FLAG was evaluated with an anti-Flag antibody (left panel) or Proteintech anti-TANGO2 antibody. The green dashed square highlight bands detected with the two antibodies, indicating their specificity. The red dotted lines highlight the unspecific band characteristic of the Proteintech antibody between 70 and 100 kDa. J TANGO2 C-ter WT forms an oligomer between 100-130 kDa. K Beta-oxidation flux in whole blood of TANGO2 patients compared to controls. No abnormalities of beta-oxidation flux were observed in patients compared to controls. No difference or **p<0,01 from unpaired Student t-tests.

### Slide 3
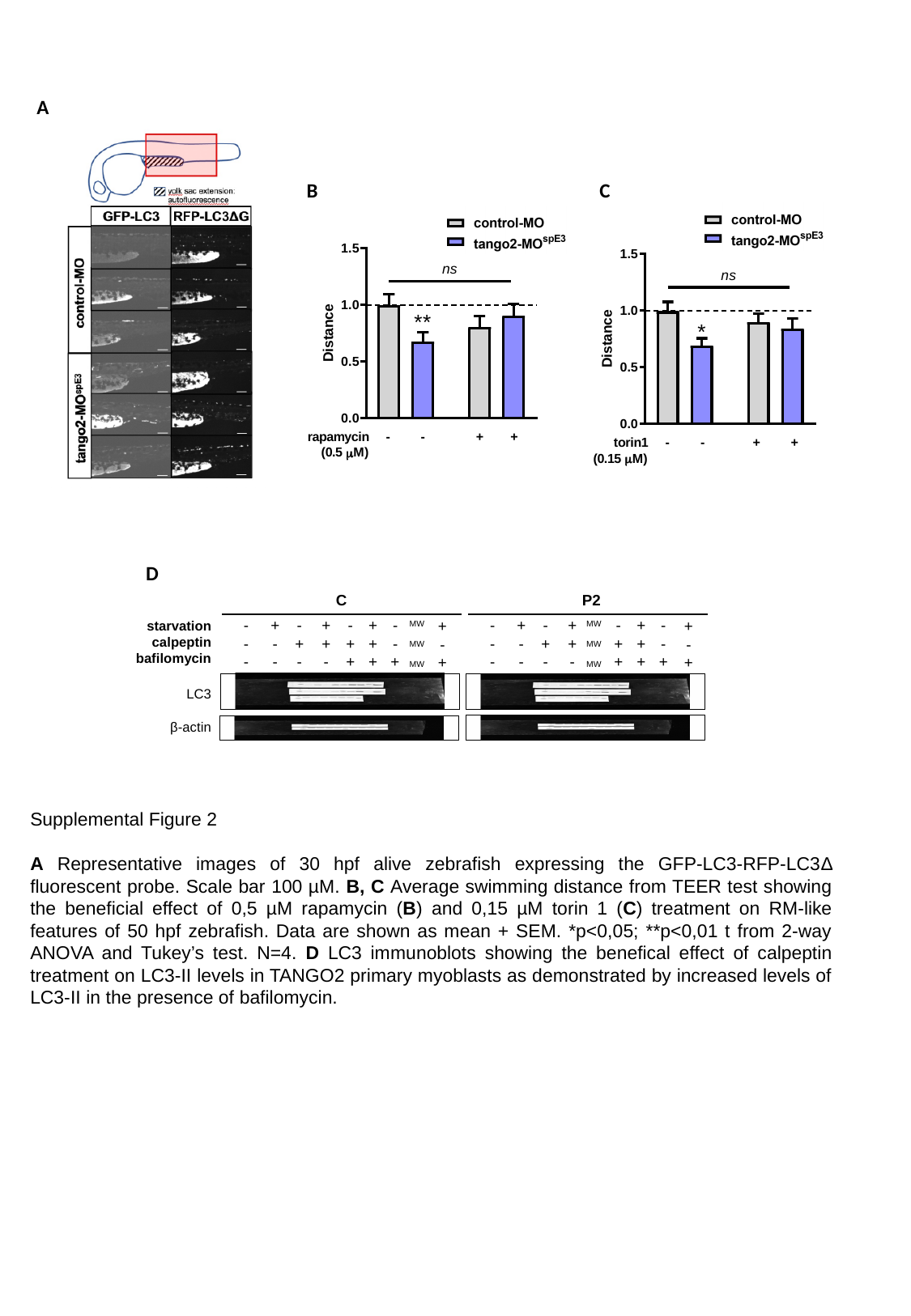

A
B
C
D
C
P2
-
-
-
+
-
-
-
+
-
+
+
-
-
+
+
+
+
+
-
-
+
-
-
-
+
-
-
-
+
-
+
+
-
-
+
+
+
+
+
-
-
+
+
-
+
+
-
+
starvation
calpeptin
bafilomycin
MW
MW
MW
MW
MW
MW
LC3
β-actin
Supplemental Figure 2
A Representative images of 30 hpf alive zebrafish expressing the GFP-LC3-RFP-LC3Δ fluorescent probe. Scale bar 100 µM. B, C Average swimming distance from TEER test showing the beneficial effect of 0,5 µM rapamycin (B) and 0,15 µM torin 1 (C) treatment on RM-like features of 50 hpf zebrafish. Data are shown as mean + SEM. *p<0,05; **p<0,01 t from 2-way ANOVA and Tukey’s test. N=4. D LC3 immunoblots showing the benefical effect of calpeptin treatment on LC3-II levels in TANGO2 primary myoblasts as demonstrated by increased levels of LC3-II in the presence of bafilomycin.
